# Evolutionary rescue in a fluctuating environment

**DOI:** 10.1101/2022.11.03.515107

**Authors:** Loïc Marrec, Claudia Bank

## Abstract

No environment is constant over time, and environmental fluctuations impact the outcome of evolutionary dynamics. Survival of a population not adapted to some environmental conditions is threatened unless a mutation rescues it, an eco-evolutionary process termed evolutionary rescue. We here investigate evolutionary rescue in an environment that fluctuates between a favorable state, in which the population grows, and a harsh state, in which the population declines. We develop a stochastic model that includes both population dynamics and genetics. We derive analytical predictions for the mean extinction time of a non-adapted population given that it is not rescued, the probability of rescue by a generalist mutation, and the mean appearance time of a rescue mutant, which we validate using numerical simulations. We find that evolutionary rescue is less (respectively more) likely if the environmental fluctuations are stochastic rather than deterministic and if the mean time between each environmental change is less (respectively more) than the mean survival time of the population in the harsh environment. We demonstrate that high equilibrium population sizes and *per capita* growth rates maximize the chances of evolutionary rescue. We show that an imperfectly harsh environment, which does not fully prevent births but makes the death rate to birth rate ratio much greater than unity, has almost the same rescue probability as a perfectly harsh environment, which fully prevents births. Finally, we put our results in the context of antimicrobial resistance and conservation biology.

## 1 Introduction

Environmental change happens all around us and impacts the populations that experience it. For example, every living organism is exposed to climate change [1, 2, 3, 4], and pathogenic microbes are treated with varying drug concentrations [5, 6], which threatens their survival. Populations too poorly adapted to changing environmental conditions may go extinct unless adaptive mutations counteract their decline, a process termed evolutionary rescue. An important question in theoretical biology is to predict whether evolutionary rescue will occur before extinction and which conditions favor adaptation [7, 8, 9].

Numerous theoretical works have shown that environmental fluctuations affect evolutionary dynamics. Specifically, analytical predictions were derived to assess the fate of a mutation in a fluctuating environment, which impacts either demography [10, 11, 12] or selection [12, 13, 14, 15]. For example, these analytical predictions showed that a cyclic change in population size or selection coefficient (resembling a fluctuating environment) results in a mutant fixation probability that is also periodic as a function of the time of appearance. However, many of these models assume that environmental fluctuations do not impact demography and selection together, which is a simplification that overlooks a key aspect of evolutionary dynamics: the interaction of population dynamics and population genetics (but see [16]).

The interaction between population dynamics and genetics is all the more important as it exists everywhere in nature. For example, antimicrobial treatments are designed to decrease the size of microbial populations until their eradication, which inhibits reproduction and thus the appearance of mutation, but selects for antimicrobial resistant mutants that may appear during drug therapy [5, 17, 18, 19, 20, 21, 22, 23, 24]. Similarly, climate change may cause extinction [25, 26, 27], but some animal species adapt quickly to stressful conditions and reverse their decline [28, 29, 30]. Importantly, the interaction between demography and selection can result in population decline, reducing genetic diversity, which could facilitate evolutionary rescue [31]. To improve theoretical predictions and inference from experimental and empirical data, there is a need for mathematical models that make an explicit link between ecology, evolution, and demography when quantifying the fate of a population evolving in a fluctuating environment [32, 33]. One of the challenges to overcome is to go beyond the approximation that environmental and evolutionary time scales are decoupled [34, 35]. Specifically, environmental effects are often self-averaged if environmental fluctuations are rapid [36], or a constant environment is assumed if environmental fluctuations are slow [37] (but see [11, 14, 38]). Another challenge is to derive exact analytical predictions that do not rely on deterministic or diffusion approximations, which have been shown to poorly describe extreme events such as extinction [39], yet necessary for modeling evolutionary rescue.

In this paper, we develop a minimal model that integrates population dynamics and genetics to quantify evolutionary rescue in a fluctuating environment. Specifically, we study a haploid population evolving in an environment fluctuating between a favorable state, in which the population grows, and a harsh state, in which it declines. The population is initially monomorphic, and mutants can appear upon reproduction. If a mutation unaffected by environmental changes becomes fixed, the population is rescued from extinction. Importantly, we investigate the probability of evolutionary rescue using a stochastic framework with numerical and analytical tools, resulting in an exact computation of the population’s fate under deterministic versus stochastic environmental fluctuations. We compare a perfectly harsh (i.e., fully birth-preventing) and an imperfectly harsh (i.e., not fully birth-preventing) environment and identify which growth parameters promote evolutionary rescue using different growth types.

## 2 Model and methods

### A population model in a fluctuating environment

We study a wild-type population of size *N*_*W*_, which can vary over time and is limited by a carrying capacity *K*. Each wild-type individual has the same birth rate *b*_*W,α*_, which depends on the environmental state, and death rate *d*_*W*_. The population follows a logistic growth in which the *per capita* birth rate satisfies *b*_*W,α*_(1−*N*_*W*_ */K*), and the *per capita* death rate is equal to the intrinsic death rate. We also present results for the Gompertz and Richards growths, whose *per capita* birth rates satisfy *b*_*W,α*_ log(*K/N*_*W*_) and *b*_*W,α*_(1− (*N*_*W*_ */K*)^*β*^), respectively (see figure 1d). These growth types, which are used to fit population growth data [40, 41], have different equilibrium sizes and *per capita* growth rates that may impact the probability of evolutionary rescue. The population evolves in an environment that fluctuates between two states, namely favorable F and harsh H, which impacts only the birth rate. In the favorable environment, the *per capita* birth rate is larger than the death rate (e.g., *b*_*W,F*_ (1−*N*_*W*_ */K*) *> d*_*W*_ for the logistic growth) so that the population grows towards its equilibrium size 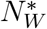. Conversely, in the harsh environment, the *per capita* birth rate is lower than the death rate (e.g., *b*_*W,H*_(1−*N*_*W*_ */K*) *< d*_*W*_ for the logistic growth), so that the population declines towards extinction. An example of a simulation run is shown in figure 1a). The environment remains in each state for a duration *τ*, sampled from the probability density function ℱ_*τ*_. In the case of deterministic fluctuations, we set ℱ_*τ*_ (*t*) = *δ*(*t*−*τ*), in which *δ* is the Dirac delta. In the case of stochastic fluctuations, the phase duration is drawn from a biased normal distributions of mean *τ* and standard deviation *σ* given by 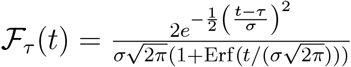 (see figure 1b-c) that exclude negative values.

**Figure 1:**
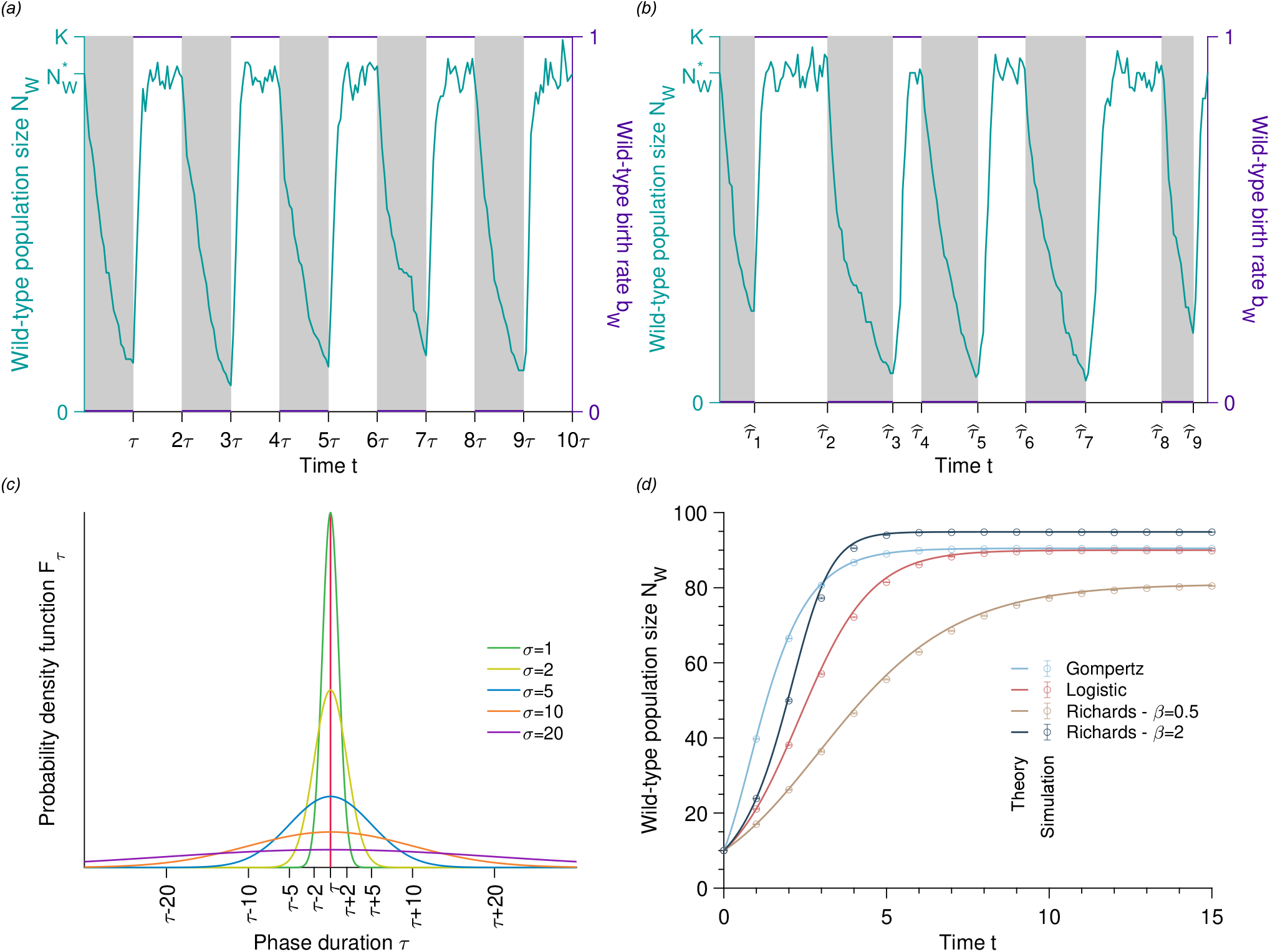
Illustration of the model - Population dynamics in a fluctuating environment. Wild-type population size and birth rate versus time with deterministic *(a)* and stochastic switches *(b)*. In both panels, the solid line represents a realization of a simulation run under the logistic growth. The gray and white phases correspond to harsh and favorable environments, respectively. *(c)* Probability density function of the phase duration, which is normally distributed, positive, of mean *τ* and standard deviation *σ. (d)* Population size versus time for different population growth patterns in a constant favorable environment. Solid lines represent analytical predictions, and data points show simulated data averaged over 10^4^ stochastic realizations. Error bars correspond to the 95% confidence intervals. Parameter values: wild-type birth rate in favorable environment *b*_*W,F*_ = 1, wild-type birth rate in harsh environment *b*_*W,H*_ = 0, wild-type death rate *d*_*W*_ = 0.1, carrying capacity *K* = 100, and equilibrium wild-type population size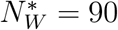.

A generalist mutant appears upon reproduction with probability *μ* and has birth *b*_*M*_ and death rates *d*_*M*_ constant across environments. We assume that the population is initially monomorphic for the wild type and that its initial population size equals the equilibrium size 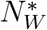. Demographic equilibrium, obtained when births and deaths offset each other, is equal to *K*(1 − *d*_*W*_ */b*_*W,F*_) for the logistic growth. Our analytical approach uses methods from birth-death processes described by master equations [42, 43]. Our simulations are based on a Gillespie algorithm and incorporate individual stochastic division, mutation, and death events [44, 45]. The algorithm we used is detailed in the Supplemental Material.

### Timescales of environmental fluctuations

In the fluctuating environment, either the population goes extinct at time *T*_0_, or a mutant appears, fixes, and thus rescues the population before *T*_0_. The evolutionary outcome crucially depends on how the environmental timescale *τ* compares to the population’s lifetime *τ*_0,*H*_ in the harsh environment. In the limit of large *τ*, for *τ* ≫*τ*_0,*H*_, very slow environmental fluctuations lead to rapid extinction (i.e., *T*_0_ = *τ*_0,*H*_) because the harsh environment lasts much longer than the population lifetime in the harsh environment (see figure S1a). Here, the rapid extinction leaves little (if *b*_*W,H*_ *>* 0 and *d*_*W*_ */b*_*W,H*_ ≫1) or no opportunity (if *b*_*W,H*_ = 0) for rescue mutants to appear and therefore the rescue probability *p*_*r*_ is likely to be zero. In the limit of small *τ*, for *τ* ≪*τ*_0,*H*_, very rapid environmental fluctuations make the population persist long enough for mutations to arise and rescue it. In the particular case of very fast environmental fluctuations, the evolutionary dynamics can be described by a constant environment with an averaged birth rate 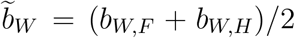 and an effective equilibrium size 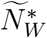 satisfying 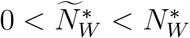 (see figure S1b). Although rapid environmental fluctuations maintain the population in an equilibrium state, its extinction time 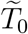 is reduced compared to if it remained indefinitely in the favorable environment. In the case of an effective constant environment, the mean appearance time of a beneficial mutant of selection coefficient 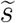 (i.e., 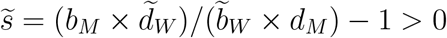) that becomes fixed is given by 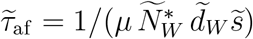 (see figure S2). If this time is much shorter than the mean extinction time 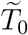, the rescue time probability *p*_*r*_ is likely to be one.

In the following, we focus on nontrivial cases in which the environmental timescale is of the same order of magnitude as the population lifetime in the harsh phase (i.e., *τ* ∼ *τ*_0,*H*_) and the rescue probability is likely to satisfy 0 *< p*_*r*_ < 1.

### Stochastic dynamics of the wild-type population

We describe the population dynamics as a Markovian birth-death process that includes stochasticity inherent to demographic noise [42, 43]. More specifically, the probability that a population has a given size between 0 and *K* at a given time *t* is described by a system of *K* +1 differential equations. This system is coupled since a population jumps from one to another size with a rate depending on its current size. The system of differential equations, called the master equation, governs the time-evolution of the probability *P*_*α*_(*N*_*W*_, *t* | *N*_*W*,0_) of having *N*_*W*_ individuals at time *t* in the environmental state *α* given that *N*_*W*,0_ were initially present, and reads for the logistic growth

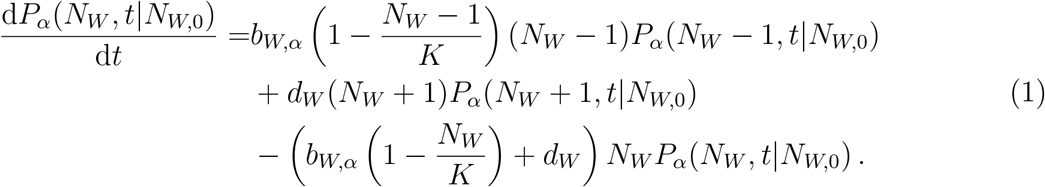

We write equation (1) in a matrix form, 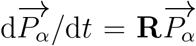 where 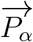 is the probability column vector and **R** the *K* + 1 × *K* + 1 transition rate matrix. The solution of equation (1) reads 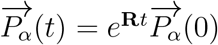 where 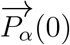 is the initial condition column vector, whose 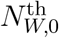 row is equal to 1 whereas the others are zero. We assume that each favorable phase is long enough for the population to reach its equilibrium size, that is *τ* ≫ 1*/*(*b*_*W,F*_ − *d*_*W*_), given that the population is not extinct. Then, each harsh phase starts from a population size 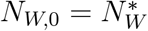. When the harsh environment fully prevents the birth of individuals (i.e., *b*_*W,H*_ = 0), equation (1) is analytically solvable, and we obtain

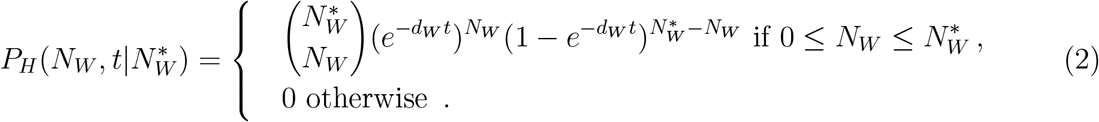

The previous equation shows that the population size *N*_*W*_ is sampled according to a normal distribution 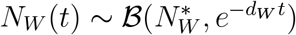 in the harsh environment. When the harsh environment does not fully prevent the birth of individuals (i.e., *b*_*W,H*_ ≠ 0), equation (1) is solved numerically. To obtain the wild-type population size at any time *t* given that it has not gone extinct, we need to modify the master equation since equation (1) includes the stochastic trajectories leading to rapid extinction [39]. To exclude these trajectories, we consider a biased master equation giving the probability *Q*_*α*_(*N*_*W*_, *t*|*N*_*W*,0_) = *P*_*α*_(*N*_*W*_, *t* | *N*_*W,α*_)*/*(1 − *P*_*α*_(0, − *t* |*N*_*W*,0_)) of having *N*_*W*_ individuals at time *t* in the environmental state *α*, given that *N*_*W*,0_ were initially present and that the population is not extinct [12, 46]. This master equation is similar to equation (1) with an additional term *Q*_*α*_(*N*_*W*_, *t* | *N*_*W*,0_)(d*P*_*α*_(0, *t* |*N*_*W*,0_)*/*d*t*)*/*(1 −*P*_*α*_(0, *t* |*N*_*W*,0_)). Note that *P*_*F*_ (0, *t* |*N*_*W*,0_) can be analytically obtained by linearizing the master equation, whereas *P*_*H*_(0, *t* |*N*_*W*,0_) is computed numerically if *b*_*W,H*_ *>* 0, or using equation (2) if *b*_*W,H*_ = 0. The mean population size in the environmental phase *α* starting from *N*_*W*,0_ individuals is given by 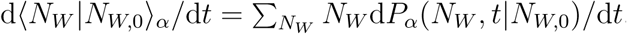 which for the logistic growth leads to

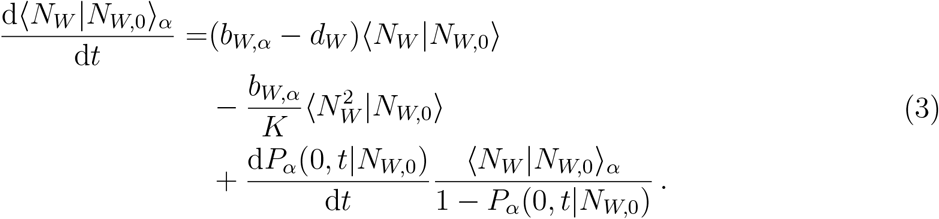

The initial condition *N*_*W*,0_ is equal to 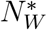 in the harsh environment, whereas it is more difficult to obtain it in the favorable environment. That is because the initial size in the favorable phase is random and depends on the previous harsh phase. Thus, we calculate the mean population size in the favorable phase by summing the trajectories with all possible initial conditions *N*_*W*,0_ weighted by their respective probability

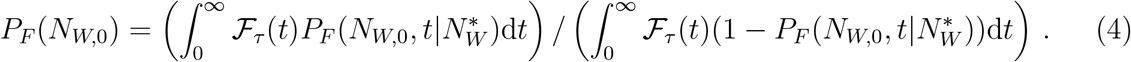

In other words, *P*_*F*_ (*N*_*W*,0_) is the probability of having *N*_*W*,0_ individuals at the end of each harsh phase, given that the population is not extinct. The denominator of equation (4) ensures that only trajectories in which the population did not go extinct in the previous harsh phase are considered. This yields

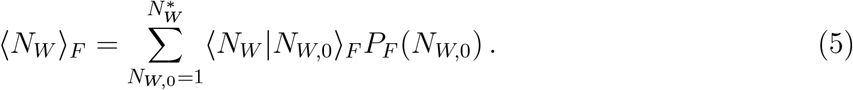

Now that we have quantified the dynamics in both environments, namely favorable and harsh, we can write the complete dynamics as

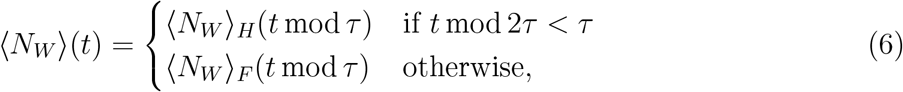

where *m* mod *n* is the modulo operation that yields the remainder of the division of *m* by *n*. An example in which equation (6) is tested against simulated data is shown in figure S3. A technical point should be made clear. Since our birth rates are nonlinear, unless *b*_*W,H*_ = 0, the mean population size given by equation (3) depends on higher-order moments. In other words, the system of equations for the moment dynamics is not closed. Because the system of moment equations is not closed, we apply a binomial moment closure approximation as it proves to be the best for the logistic growth. In contrast, we use a mean field approximation (equivalent to a deterministic equation) for the Gompertz and Richards growths (citation to come).

### Appearance and fixation of a mutant

A beneficial mutant rescues a population from extinction only if it survives the initial drift phase at low frequencies and becomes fixed. Following [12, 47], the fixation probability reads

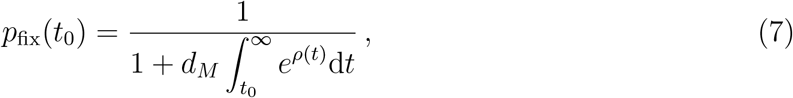

where *t*_0_ is the appearance time and

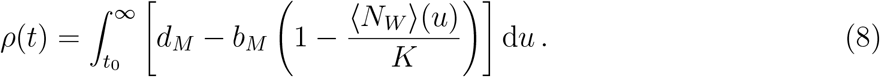

The previous equation applies to the logistic growth. To obtain the fixation probability for the Gompertz and Richards growths, *b*_*M*_ must be multiplied by log(*K/* ⟨*N*_*W*_⟩) and (1 − (⟨*N*_*W*_⟩ */K*)^*β*^), respectively. Equation (7) shows that the fixation probability of the mutant in our eco-evolutionary model depends on its appearance time (see figure S4). In contrast, the fixation probability is constant for fixed and infinite population sizes in a constant environment.

In our model with fluctuating selection coefficients, a mutant is more likely to fix in the harsh phase than in the favorable phase since *d*_*W*_ */b*_*W,F*_ *> d*_*W*_ */b*_*W,H*_. If the harsh environment fully prevents the reproduction of wild-type individuals, the fixation probability of a mutant is maximal just before the beginning of the harsh phase [47, 48, 49].

Since environmental fluctuations lead to varying birth rates, the number of mutants that appear per unit of time is not constant. We calculate the probability that a mutant appears and fixes between 0 and *t* given that none has done so before as *p*_af_(*t*) = 1 − *e*^*−*Σ(*t*)^ [47], where

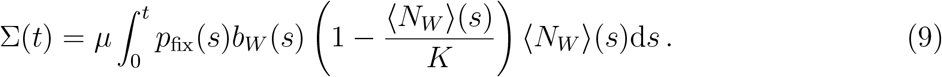

Here, the wild-type birth rate depends on time because of the assumed environmental fluctuations. Taking the limit *t* → ∞ of *p*_af_(*t*) yields the rescue probability *p*_*r*_. Note that equation (9) applies to the logistic growth model. To adjust equation (9) for the Gompertz and Richards growths, *b*_*W*_ (*s*) must be multiplied by log(*K/*⟨*N*_*W*_ ⟩) and (1 − (⟨*N*_*W*_ ⟩*/K*)^*β*^), respectively.

### Data availability

Simulations were performed with C (version gcc-9) and Matlab (version R2021a). All annotated code to repeat the simulations and visualizations is available at https://github.com/LcMrc and will be deposited on Zenodo upon acceptance of the paper.

## 3 Heuristic analysis

### Two different extinction mechanisms contribute to failed evolutionary rescue

Environmental fluctuations decrease the persistence time of a population if they induce paths to extinction. In the harsh environment, the population declines because the death rate exceeds the birth rate. If the harsh phase duration *τ* is longer than the survival time *τ*_0,*H*_, the population goes extinct. The survival time in the harsh environment is stochastic and extinction occurs with probability 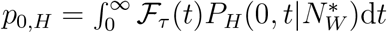 (see equation (2)).

A second path to extinction exists in the favorable environment. If the population survives the previous harsh phase, it possibly starts the new favorable phase with few individuals. Small initial population sizes lead to strong demographic noise that may drive the population to extinction with probability 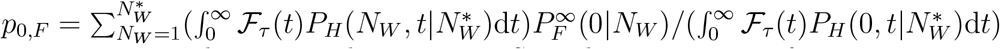 [39]. An example of each extinction mechanism is shown in Fig S5. The proportion of extinctions occurring in the favorable phase, which is given by *ω*_*F ≈*_ *p*_0,*F*_ */*(*p*_0,*F*_ + *p*_0,*H*_), is expected to decrease with increasing phase duration. The longer the harsh phase, the more certain it drives the population to extinction. Conversely, short harsh phases do not drive the population to extinction but, in some cases, decrease the population size enough to lead to rapid extinction in the next favorable phase.

### Stochastic environmental switches can increase or decrease the rescue probability

As explained before, the fate of the population depends on how the survival time of the wild-type in the harsh environment compares to the phase duration. If the environmental fluctuations are deterministic and the mean phase duration is shorter than the mean survival time in the harsh environment (i.e., *τ < τ*_0,*H*_), the harsh phase is too short to drive the population to extinction. However, if the environmental fluctuations are stochastic, some harsh phases are longer than average, which favors extinction and decreases the total extinction time and the rescue probability. If the environmental fluctuations are deterministic and the mean phase duration is longer than the mean survival time in the harsh environment (i.e., *τ*_0,*H*_ *< τ*), the harsh phase is long enough to drive the population to extinction. However, if the environmental fluctuations are stochastic, some harsh phases are shorter than average, which favors population survival and increases the total extinction time and the rescue probability. In summary, no matter whether the environmental fluctuations are deterministic or stochastic, the total extinction time and the probability of rescue decrease as the phase duration increases. However, for a mean phase duration shorter than the mean survival time in the harsh environment (i.e., *τ < τ*_0,*H*_), the stronger the environmental stochasticity, the lower the total extinction and the rescue probability. The opposite is valid for a mean phase duration longer than the mean survival time in the harsh environment (i.e., *τ*_0,*H*_ *< τ*).

### Small birth rates in the harsh environment leave rescue probabilities almost unchanged

The harsh environment induces a wild-type birth rate lower than the death rate. Specifically, a perfectly harsh environment fully prevents births, whereas an imperfectly harsh environment allows for a small number of births during the harsh phase. As long as the birth rate in the harsh environment is much lower than the death rate (i.e., *d*_*W*_ */b*_*W,H*_ ≫ 1; e.g., *d*_*W*_ */b*_*W,H*_ = 10), the population is driven to extinction on a time scale equal to 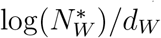 (see figure S6). Thus, the mean total extinction time should not significantly differ between perfectly and imperfectly harsh environments that satisfy *d*_*W*_ */b*_*W,H*_ ≫ 1. However, *b*_*W,H*_ may impact the rescue probability as it determines how many births occur and how many mutants appear. Specifically, there are births in each 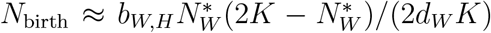 harsh phase, and the probability that at least one mutant appears is given by 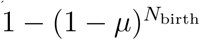 (see figure S7). Thus, the larger the birth rate *b*_*W,H*_ and the mutation probability *μ*, the more mutants appear in the harsh environment. However, if the death rate to birth rate ratio satisfies *d*_*W*_ */b*_*W,H*_ ≫ 1, the number of births in the harsh environment is expected to be very small, and the number of mutants that appear is much smaller. Therefore, the rescue probability in an imperfectly harsh environment is likely similar to that in a perfectly harsh environment.

### Rescue probability depends on population growth types

In addition to studying evolutionary rescue under the logistic growth, we also present results for the Gompertz and Richards growths. Each of these growth types has a different equilibrium size 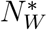 and growth rate, which affect the total extinction time and the rescue probability. First, the larger the equilibrium size, the longer it takes for the population to go extinct in the harsh environment since 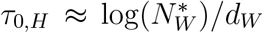 Thus, the probability *p*_0,*H*_ of extinction in a harsh environment at a given phase duration *τ* decreases as the equilibrium size increases. Second, the faster the growth, the lower the demographic stochasticity. The probability of rapid extinction *p*_0,*F*_ of a population with initial size *N*_*W*,0_ is very small compared to its equilibrium size, which is given by 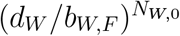 for the logistic and Richards growths and 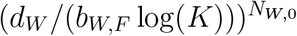 for the Gompertz growth. Thus, the probability of extinction *p*_0,*F*_ for a given phase duration decreases for populations with a higher growth rate. An increased total extinction time leaves more opportunities for mutants to appear and become fixed. Moreover, a larger growth rate results in more births, resulting in more mutants and, therefore, a higher rescue probability. As a result, growth according to the Gompertz and Richards growths with *β >* 1 is likely to favor evolutionary rescue over the logistic and Richards growths with *β <* 1 (see figure 1a).

## 4 Formal analysis

### Extinction time

From the extinction probabilities, namely *p*_0,*F*_ and *p*_0,*H*_ (see Model and methods), we compute the probability *P*_*qF*_ that the population undergoes 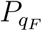 favorable phases before it goes extinct as

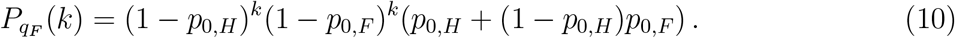

The favorable phases in which a rapid extinction occurs are excluded from this count because we focus on the favorable phases in which a mutant is most likely to appear. We obtain the mean number of favorable phases before extinction by calculating 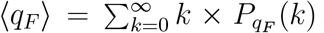, which yields

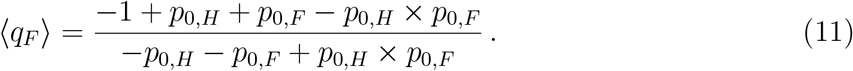

Equation (11) shows that both extinction mechanisms (extinction in the harsh phase vs. extinction due to low numbers at the beginning of the favorable phase) are important in assessing population persistence. The probabilities *p*_0,*F*_ and *p*_0,*H*_ increase as the phase duration increases (see figure S8a-b), reducing ⟨*q*_*F*_⟩. Specifically, the probability of extinction in the harsh environment ranges from 0 to 1 (i.e., 0 *< p*_0,*H*_ < 1) since short phases do not leave enough time for the population to go extinct. In contrast, long phases surely drive it to extinction before the next environmental change. The probability of extinction in the favorable environment ranges from 0 to *d*_*W*_ */b*_*W,H*_ (i.e., 0 *< p*_0,*F*_ *< d*_*W*_ */b*_*W,H*_), where *d*_*W*_ */b*_*W,H*_ is equal to the probability that a population starting with one individual rapidly goes to extinction. Using equation (11) and the proportion *ω*_*F*_ of extinction in the favorable environment (i.e., *ω*_*F*_ ≈ *p*_0,*F*_ */*(*p*_0,*F*_ + *p*_0,*H*_)), we derive the mean total extinction time as

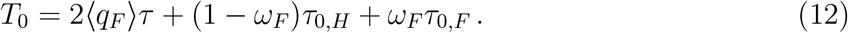

Independent of whether extinction occurs in the favorable or harsh environment, the population persists during ⟨*q*_*F*_⟩ epochs of a mean duration *τ* plus the mean survival time in the favorable (respectively harsh) environment, given that the population goes extinct, weighted by the probability that extinction occurs in the favorable (respectively harsh) environment. The mean total extinction time ranges from *τ*_0,*H*_ to 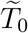 (i.e., 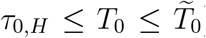), where the mean survival time *τ*_0,*H*_ in the harsh environment is obtained for very long phase durations. In contrast, the mean extinction time 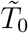 in an effectively constant environment is obtained for very short phase durations. The proportion *ω*_*F*_ of extinction in the favorable environment decreases as the phase duration increases (see figure S8c), reducing *T*_0_. If the ratio of death rate to birth rate in the favorable environment is much smaller than unity (i.e., *d*_*W*_ ≪ *b*_*W,F*_), we can assume that rapid extinction in the favorable environment occurs only if the population starts with a single individual, hence *τ*_0,*F*_ ≈ 1*/d*_*W*_. The extinction time *τ*_0,*H*_ in the harsh environment is then approximately equal to log 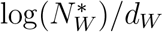 if *τ* > *τ*_0,*H*_, or to *τ* otherwise. Our analytical predictions accurately predict the simulated data (see figure 2; see also figure S9 for ⟨*q*_*F*_⟩). As reported in figure 2a), the greater the environmental stochasticity (i.e., the larger the standard deviation *σ*), the smaller the mean total extinction time *T*_0_. Even small values of the standard deviation of environmental stochasticity *σ* dramatically affect population persistence. The mean total extinction time for deterministic and stochastic fluctuations intersect around the mean survival time in the harsh environment. Beyond this time, the population persists longer in an environment with highly stochastic fluctuations, but the difference to the result for deterministic fluctuations becomes much smaller. As described by Jensen’s theorem [50], as soon as the mean total extinction time with deterministic fluctuations resembles a convex function, addition of stochasticity reduces this convexity. As reported in figure 2b), populations growing under a growth type with a larger equilibrium size and growth rate have an increased extinction time. This difference fades as the phase duration increases since extinction occurs mainly in the harsh environment, where the extinction probability is independent of the growth type. Finally, figure 2c) shows that for any ratio *d*_*W*_ */b*_*W,H*_ much greater than unity, the mean total extinction time is equal because the probability of extinction in the harsh environment is the same as if *b*_*W,H*_ = 0. Note that the maximum population size scales the window of phase durations that lead to non-trivial rescue probabilities (i.e., *τ* ∼ *τ*_0,*H*_ so that 0 *< p*_*r*_ < 1). Specifically, the mean survival time in the harsh environment is given by 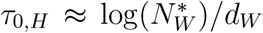. In contrast, its variance is approximately equal to 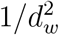 (both quantities can be derived from equation 2). We present additional results for different maximum population sizes as a function of *τ /τ*_0,*H*_ in figure S10.

**Figure 2:**
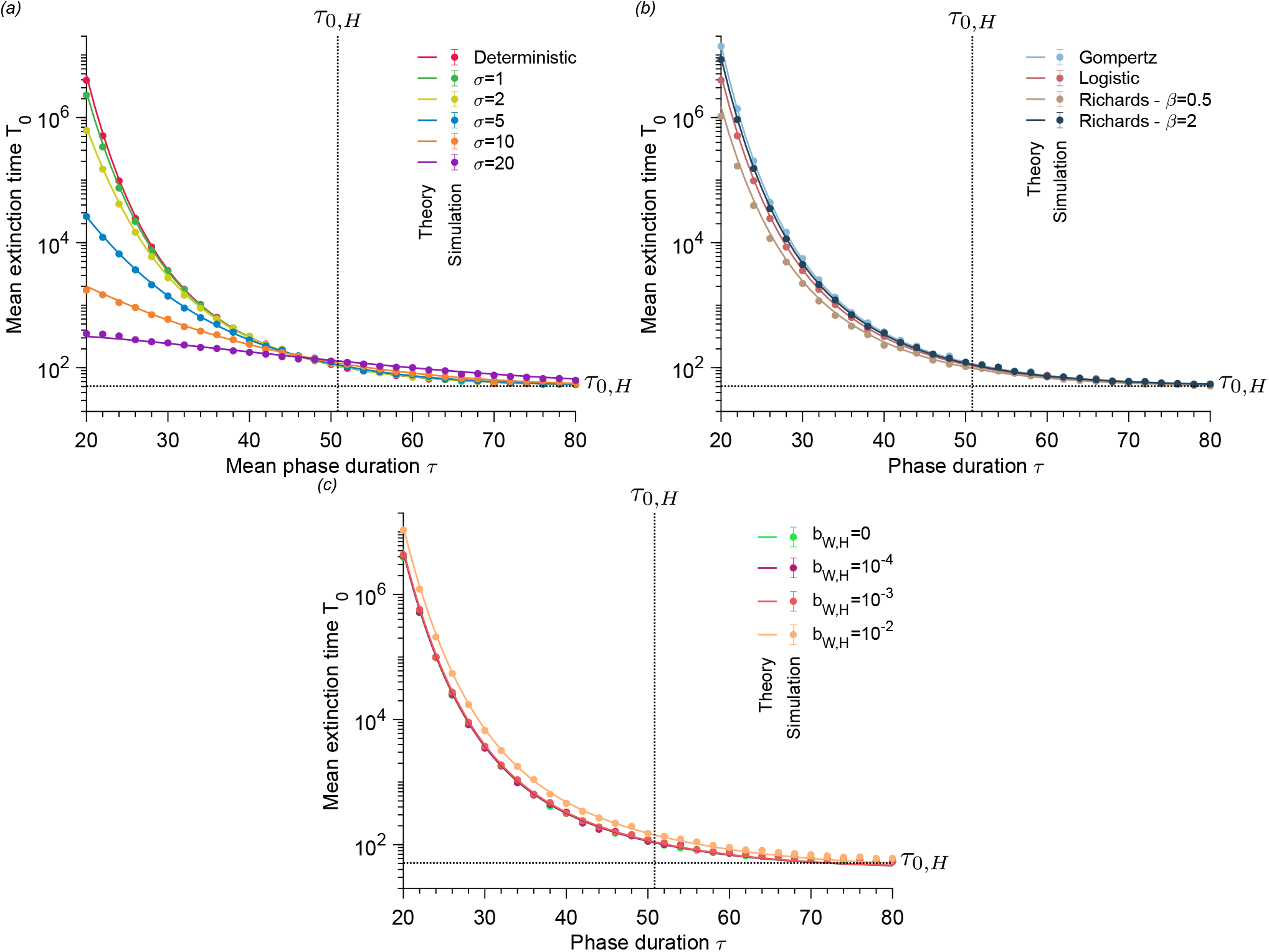
Extinction time decreases as phase duration increases. Mean extinction time versus phase duration. Panel *(a)* compares deterministic and stochastic switches for the logistic model, panel *(b)* compares different population growth models, and panel *(c)* compares a perfectly harsh environment to imperfectly harsh environments for the logistic model. The solid lines represent analytical predictions, and the points simulated data averaged over 10^4^ stochastic realizations. The error bars correspond to the 95% confidence intervals. Vertical dotted lines represent the mean survival time in the harsh environment. Parameter values: wild-type birth rate in favorable environment *b*_*W,F*_ = 1, wild-type birth rate in harsh environment *b*_*W,H*_ = 0 (in *a* and *b*), wild-type death rate *d*_*W*_ = 0.1, carrying capacity *K* = 100, and equilibrium wild-type population size 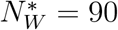.

### Rescue probability

Using the mean number ⟨*q*_*F*_⟩ of favorable phases that the population undergoes, we calculate the probability that a generalist mutant (i.e., one not affected by environmental fluctuations) appears and takes over the population before extinction occurs. We obtain

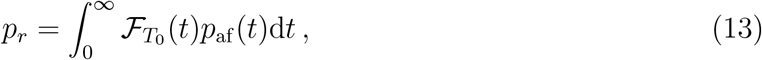

where 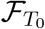 is the probability density function of the total extinction time. Equation (13) is simplified by separating the contribution of the favorable and harsh environments. Either the mutant appears in the favorable environment while the population is growing or in the harsh environment if the division is not fully hindered. Thus, the rescue probability *p*_*r*_ reads

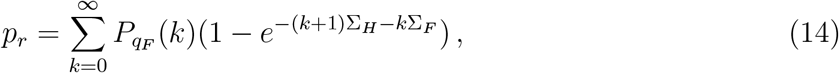

where

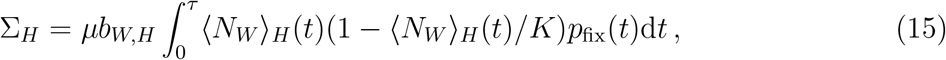

and

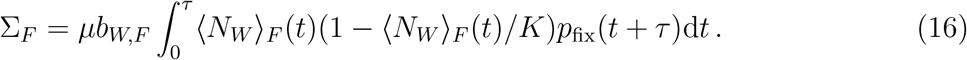

Our analytical predictions match the simulated data very well (see figure 3). In particular, figure 3a) shows that equation (14) is valid from the rare to the frequent mutation regime. All panels highlight the transition from the regime of fast fluctuations, in which *p*_*r*_ ≈ 1, to slow fluctuations, in which *p* _*r*_ ≈ 0. This transition is more abrupt for rare mutations than for frequent mutations. The more mutants there are, the more likely one mutant becomes fixed and rescues the population before extinction, hence the higher rescue probability at a given phase duration. As reported in figure 3b), environmental stochasticity decreases the chances of evolutionary rescue because it also decreases the mean total extinction time. Population growth types with the highest growth rates and equilibrium sizes have the highest rescue probabilities at a given phase duration because they lead to more mutant appearances per unit of time (see figure 3c). As shown in figure 3d), a harsh environment that does not fully prevent the reproduction of individuals leaves more opportunities for mutants to appear, resulting in a higher rescue probability.

**Figure 3:**
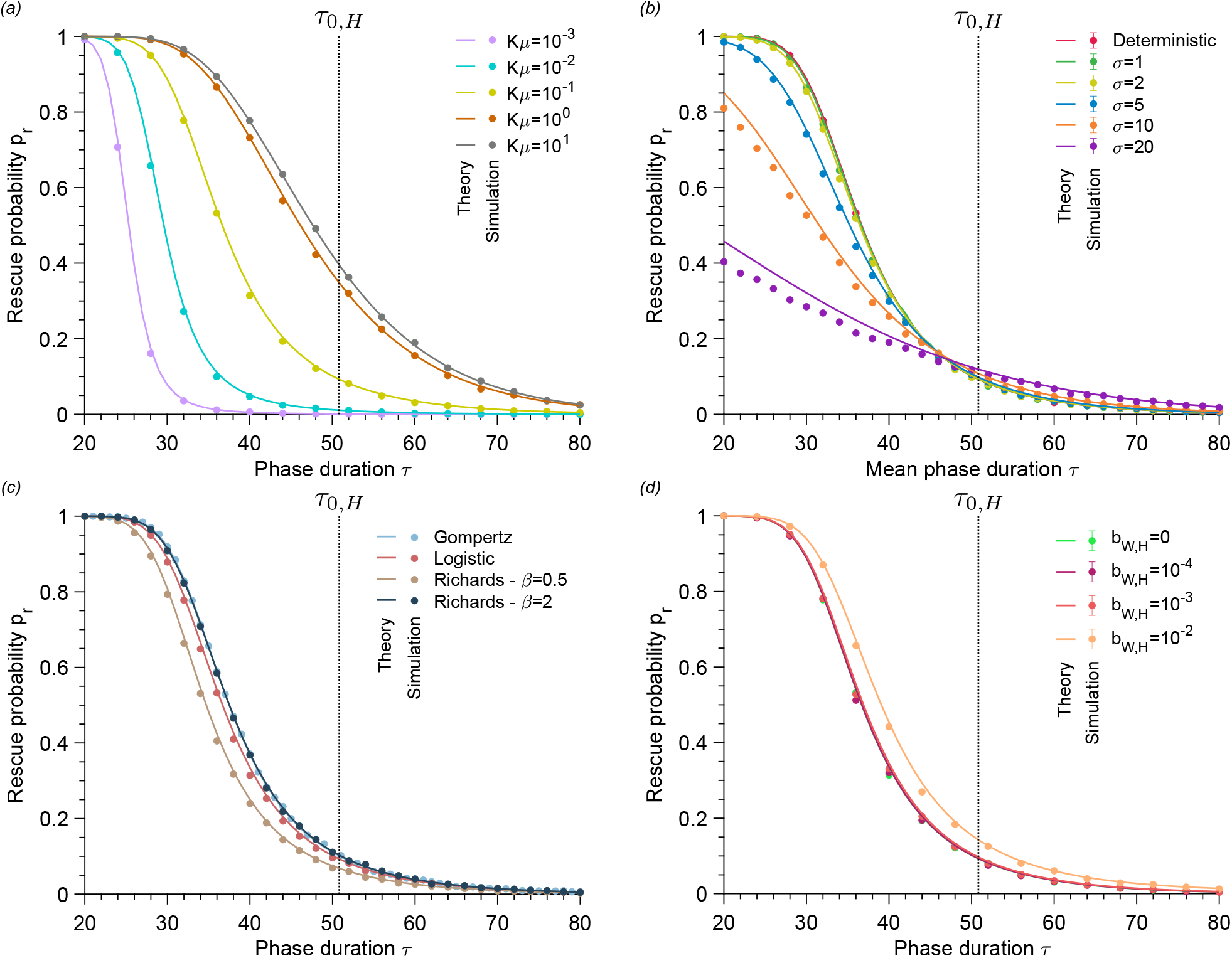
Rescue probability decreases as phase duration increases. Rescue probability versus phase duration. Panel *(a)* compares different mutation rates, panel *(b)* deterministic and stochastic switches for the logistic growth, panel *(c)* different population growth types, and panel *(d)* a perfectly vs. imperfectly harsh environment for the logistic growth. The solid lines represent analytical predictions, and the points simulated data averaged over 10^4^ stochastic realizations. The error bars correspond to the 95% confidence intervals. Vertical dotted lines represent the mean survival time in the harsh environment. Parameter values: wild-type birth rate in favorable environment *b*_*W,F*_ = 1, wild-type birth rate in harsh environment *b*_*W,H*_ = 0 (in *a, b* and *c*), wild-type death rate *d*_*W*_ = 0.1, mutant birth rate *b*_*M*_ = 1, mutant death rate *d*_*M*_ = 0.1, carrying capacity *K* = 100, mutation rate *μ* = 10^*−*3^ (in *b, c*, and *d*) and equilibrium wild-type population size 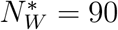.

The rescue probability is independent of the carrying capacity *K* at a given normalized phase duration *τ /τ*_0,*H*_ if the mutational influx *Kμ* is constant (see figure S11). The carrying capacity value determines the phase duration window in which the rescue probability transitions from 1 to 0 through the mean survival time in harsh environment. The product *Kμ* determines the number of mutants that appear per unit of time.

### Appearance time

We derive the average appearance time *τ*_af_ of a mutant that fixes, given that the population is rescued, as

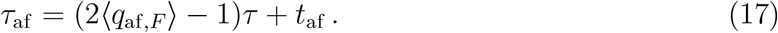

The mean number ⟨*q*_af,*F*_⟩ of favorable phases that occur before a mutant appears and fixes, given that the population is rescued, is given by

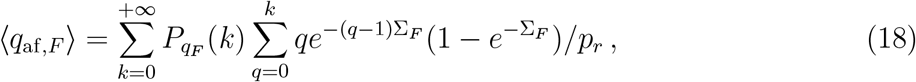

and *t*_af_ is the mean appearance time of a mutant that becomes fixed in the favorable environment. Since the mean total extinction time and rescue probability are similar for *d*_*W*_ */b*_*W,H*_ ≫ 1 (see figures 2 and 3), we assume that a mutant is unlikely to emergein the harsh environment. In the moderate to frequent mutation regime and regardless of phase duration, the mutant that rescues the population appears during the first favorable phase (see figures 4 and S12). Then *τ*_af_ increases as *τ* increases. If mutations are rare, the number of favorable phases before a rescue mutant appears decreases as the phase duration increases. More precisely, ⟨*q*_af,*F*_⟩ converges to unity when the phase duration is longer than the survival time in the harsh phase. The population goes extinct quickly for such a phase duration, so the mutant must appear in the first favorable phases. Our results confirm previous observations that the mutant rescue the population from extinction tends to appear just before an environmental change from the favorable to the harsh state [47, 48, 49].

**Figure 4:**
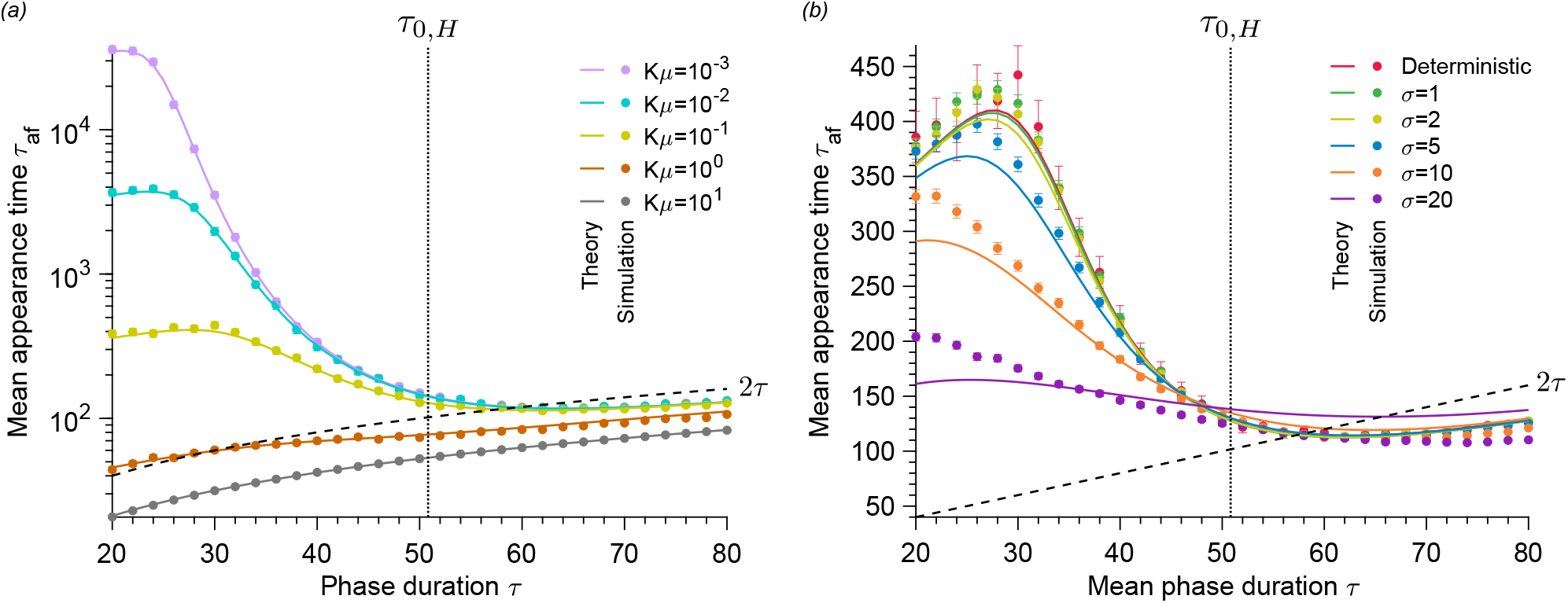
The higher the mutation rate and phase duration, the earlier the rescue mutant appears. Mean appearance time of a mutant that rescues the population versus phase duration. Panel *(a)* compares different mutation rates and panel *(b)* deterministic and stochastic fluctuations for the logistic growth. The solid lines represent analytical predictions, and the points simulated data averaged over 10^4^ stochastic realizations. The error bars correspond to the 95% confidence intervals. Vertical dotted lines represent the mean survival time in the harsh environment. Parameter values: wild-type birth rate in favorable environment *b*_*W,F*_ = 1, wild-type birth rate in harsh environment *b*_*W,H*_ = 0, wild-type death rate *d*_*W*_ = 0.1, mutant birth rate *b*_*M*_ = 1, mutant death rate *d*_*M*_ = 0.1, carrying capacity *K* = 100, mutation rate *μ* = 10^*−*3^ (in *b*) and equilibrium wild-type population size 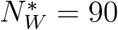.

## 5 Discussion

Whether it is microbes subjected to varying antimicrobial concentrations or animal species caught up in climate change, populations experience environmental changes threatening their survival. Determining whether populations adapt or perish is a fundamental question in many fields, from antimicrobial resistance to conservation biology. In this paper, we develop a minimal model to address evolutionary rescue in a fluctuating environment. We fully analyze our model using analytical and numerical tools from stochastic processes. Specifically, we derive equations for the extinction time, the rescue probability, and the appearance time of a rescue mutant and validate them with numerical simulations.

### Stochastic environmental fluctuations accelerate extinction and hinder evolutionary rescue compared to deterministic fluctuations

Our study quantifies the probability of evolutionary rescue of a population evolving in an environment that fluctuates, either deterministically or stochastically, between a harsh state (i.e., causing a population decline) and a favorable state (i.e., allowing population growth). We show that environmental and survival time scales determine whether stochastic environmental fluctuations favor evolutionary rescue compared to deterministic ones. Specifically, we prove that stochastic environmental fluctuations with a mean phase duration shorter than the survival time in the harsh environment dramatically decrease the mean total extinction time, the rescue probability, and the mean appearance time of a rescue mutant. Although the mean phase duration is shorter than the mean survival time, environmental stochasticity leads to longer than average phases and thus may facilitate extinction. Conversely, stochastic environmental fluctuations with a mean phase duration that is longer than the mean survival time of the population in the harsh environment very slightly increase the mean total extinction time and the rescue probability but do not significantly affect the mean appearance time of a rescue mutant.

Relating our results to a public health perspective, our model may represent treatment with biostatic drugs, which inhibit microbial division. Under this scenario, we evaluate the risk of antimicrobial resistance evolution by *de novo* mutations during therapy [8, 51]. Similar to previous theoretical work [47], we show that variation in antimicrobial concentration plays a role in the evolution of resistance. For example, we find that deterministic rapid variations favor the evolution of resistance over a constant environment. We extend [47] by showing that rapid random environmental switches of the drug concentration decrease the risk of resistance evolution. Furthermore, our analytical prediction for the probability of evolutionary rescue is valid across the regimes of fast to slow environmental fluctuations, which complements the work of [47], whose analytical results have been derived in the limit of extremely fast or slow fluctuation regimes.

Long-term therapies involving multiple dosing are subject to imperfect adherence to treatment, i.e., patients often fail to follow the exact treatment plan [52, 53, 54]. With this in mind, dose missing was theoretically investigated in [21], which showed that non-adherence allows resistant strains to grow. In our model, stochastic fluctuations may result from another form of imperfect adherence: doses taken at irregular intervals. Surprisingly, our model suggests that a biostatic antimicrobial treatment taken at irregular intervals may hinder resistance evolution rather than accelerate it.

In summary, our theoretical work can inform the design of drug treatments that prevent the evolution of resistance by choosing the best type of antimicrobial and the time interval between each dose. A possible extension would be to compare two types of antimicrobial, namely biostatic (i.e., hindering microbial division) and biocidal (i.e., killing microbes) [17, 19], by including environment-dependent death rates. We expect that biocidal drugs accelerate extinction compared to biostatic drugs while at the same time promoting evolutionary rescue. That is because since biocidal drugs do not prevent cell division, more mutants appear, which increases the probability of evolutionary rescue.

### High equilibrium population sizes and growth rates slow down extinction and favor evolutionary rescue

Our model includes an explicit link between ecology, evolution, and demography: environmental fluctuations impact the wild-type birth rate, affecting the population size and the selective advantage of the mutant. Thus, our work does not rely on the common assumption that ecology and evolution are uncoupled when studying the genetics of adaptation [55]. This assumption was already relieved in theoretical studies that have analytically predicted adaptation in a fluctuating environment inducing changes in either population size or selection coefficient, but not both together [12, 14, 38]. The analysis of our eco-evolutionary model shows that the underlying growth type (i.e., the underlying growth model) plays an essential role in the population’s fate. Specifically, we show that growth types with larger equilibrium sizes lengthen the mean survival time of the population in the harsh environment, and growth types with higher growth rates decrease the probability of rapid extinction in the favorable environment. As a result, large equilibrium sizes and high growth rates make the population persist longer and therefore favor evolutionary rescue.

Many mathematical growth models have been developed to describe population demography, from the microscopic to the macroscopic scale [40]. Mathematical growth models allow, among other things, the fitting of population dynamics data [56]. However, to date, there is no universal model that best describes any data set [41]. Our work highlights that it is crucial to correctly infer the growth type from empirical data when assessing the persistence of a population undergoing environmental change. Although we focused on haploid populations, our purely ecological results, such as extinction time, apply to diploid populations. Specifically, our model can contribute to conservation biology by guiding natural population management. For example, the birds’ breeding season was shown to be impacted by climate change, resulting in stochasticity in its duration [57]. Our model, combined with an inference of the growth type of bird populations, could allow for predictions of the risk of extinction of such populations.

Possible extensions to our model carry the potential for additional applications in conservation biology. For example, by introducing environment-dependent death rates, we may be able to identify harvesting periods that should be respected to avoid the extinction of animal populations [58]. Here, environmental fluctuations that increase the death rate may represent fishing or hunting of animal species at specific periods of the year. Our model suggests that stochastically varying fishing and hunting seasons may decrease population persistence and accelerate extinction for purely population-dynamic reasons.

### No significant differences in the impact of an imperfectly harsh environment on evolutionary rescue compared to a perfectly harsh environment

Our model compares the impact of a perfectly harsh environment (i.e., one that fully prevents births) to a perfectly harsh environment (i.e., one that does not fully prevent births) on evolutionary rescue. We show no significant differences between the two harshness levels, especially for death rates much larger than birth rates. Specifically, we prove that the mean survival time, and thus the mean total extinction time, is similar for both perfectly and imperfectly harsh environments. Although some births may occur in the imperfectly harsh environment, a mutant appearance during this phase is unlikely. Thus, an imperfectly harsh environment does not significantly favor evolutionary rescue compared to a perfectly harsh one.

This result means that our analytical results apply to an extensive range of scenarios where populations are exposed to an environment that successively causes their decline and growth. In particular, our analytical predictions for the perfectly harsh case are a good approximation for the case where the environment does not fully prevent reproduction, which is likely to be the case in nature. In the perfectly harsh case, we emphasize that our analytical predictions are explicit and exact. They do not rely on a deterministic or diffusion approximation that has been shown to poorly describe extreme events such as extinctions [39], although widely used in population genetics [10].

In summary, the randomness of environmental fluctuations is essential to consider when quantifying the persistence of a population, as is its growth type. Conversely, the harshness of the environment does not significantly impact the persistence of the population as long as it induces its decline.

## Author Contributions

LM designed the study; LM performed the numerical and analytical work; LM and CB analyzed and interpreted the data; LM and CB wrote and edited the manuscript.

## Acknowledgments

The authors thank the THEE Group and Stephan Peischl at UniBe for discussion and feedback on the manuscript. CB is grateful for funding from ERC Starting Grant 804569829 (FIT2GO) and SNSF Project Grant “MiCo4Sys”.

