## Supplementary information for "Evolutionary rescue in a fluctuating environment"

#### 1 Detailed simulation methods

In this work, the evolution of the population is simulated using a Gillespie algorithm. Let us denote by $N_W$ and $N_M$ the number of wild-type and mutant individuals, respectively. The elementary events that can happen are reproduction with or without mutation and death of an individual of either type:

- Reproduction without mutation of a wild-type individual with rate $k_W^+ = b_W(1 - (N_W + N_M)/K)(1 - \mu)$, with $b_W = b_{W,H}$ and $b_W = b_{W,F}$ in the harsh and favorable environments, respectively.
- Reproduction with mutation of a wild-type microbe with rate $k_{WM} = b_W(1 - (N_W + N_M)/K)\mu$, with $b_W = b_{W,H}$ and $b_W = b_{W,F}$ in the harsh and favorable environments, respectively.
- Death of a wild-type individual with rate $k_W^- = d_W$.
- Reproduction of a mutant individual with rate $k_M^+ = b_M(1 - (N_W + N_M)/K)$.
- Death of a mutant individual with rate $k_M^- = d_M$.

The total rate of events is given by $k_{tot} = k_W^+ + k_{WM} + k_W^- + k_M^+ + k_M^-$. The simulation steps are as follows:

1. Initialization: The population starts from $N_{W,0}$ wild-type individuals and 0 mutant individual at time $t = 0$ in the harsh environment. The next time when the environment changes is stored in the variable $t_{switch}$, which is initialized at $\tau$. Note that $\tau$ is either deterministic or sampled from a biased normal distribution (see main text).
2. The time increment $\Delta t$ is sampled randomly from an exponential distribution with mean $1/k_{tot}$, and the next event that may occur is chosen randomly, proportionally to its probability $k/k_{tot}$, where $k$ is its rate. For instance, reproduction of a wild-type individual without mutation is chosen with probability $k_W^+/k_{tot}$.
3. If $t + \Delta t < t_{switch}$, time is increased to $t + \Delta t$, and the event chosen in Step 2 is executed.

4. If $t + \Delta t \geq t_{switch}$, the event chosen in Step 2 is not executed, because an environment change has to occur before. The environment change is performed: time is incremented to $t = t_{switch}$, and the reproduction rate of the wild-type individuals is switched from $b_{W,H}$ to $b_{W,F}$ or vice-versa. In addition, $t_{switch}$ is incremented to $t_{switch} + \tau$, and thus stores the next time when the environment changes.

5. We go back to Step 2 and iterate until the total number of individuals is zero ($N_W = 0$ and $N_M = 0$) or there are only mutant individuals ($N_W = 0$ and $N_M \neq 0$).

### 2 Additional figures

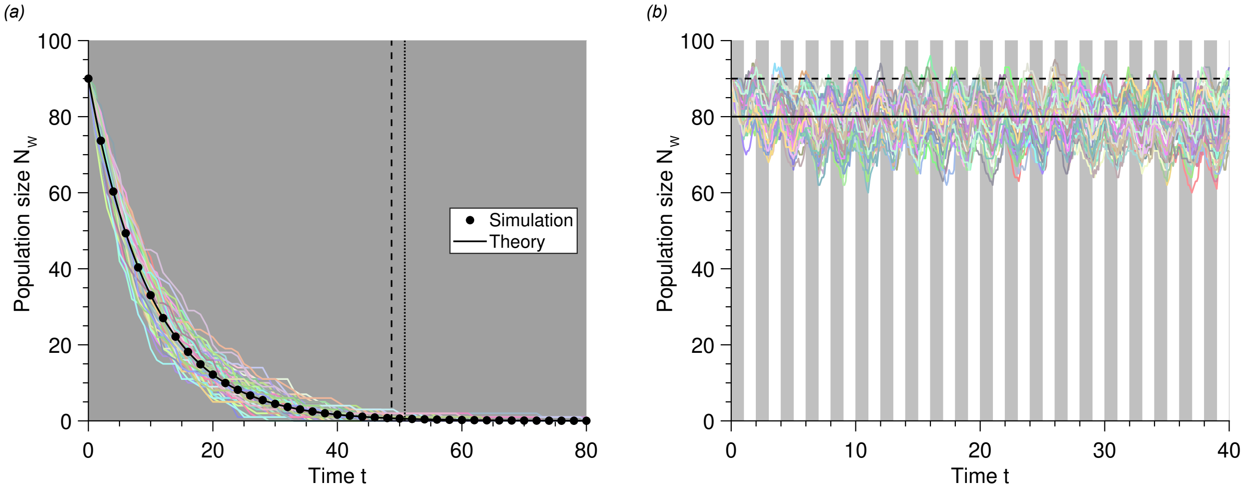


Figure 1: **Slow environmental fluctuations drive the population to extinction, whereas rapid fluctuations maintain equilibrium.** Wild-type population size $N_W$ versus time $t$ in a constant harsh environment *(a)* and fast-switching one *(b)*. In both panels, each colored line represents a realization of simulated data (the Logistic model is used in *(b)*, whereas *(a)* is valid for every growth model). The gray and white phases correspond to harsh and favorable environments, respectively. In *(a)*, the points are simulated data averaged over $10^3$ stochastic realizations, whereas the solid line corresponds to the equation $N_W(t) = N_W^* e^{-d_W t}$. The vertical dotted line represents the mean lifetime in the harsh environment. In *(b)*, the horizontal dashed, and solid lines show the equilibrium size $N_W^*$ and the averaged one $\widetilde{N_W^*}$, respectively. Parameter values: $b_{W,F} = 1$, $b_{W,H} = 0$, $d_W = 0.1$, $K = 100$, $N_W^* = 90$ and $\tau = 1$ (in $b$).

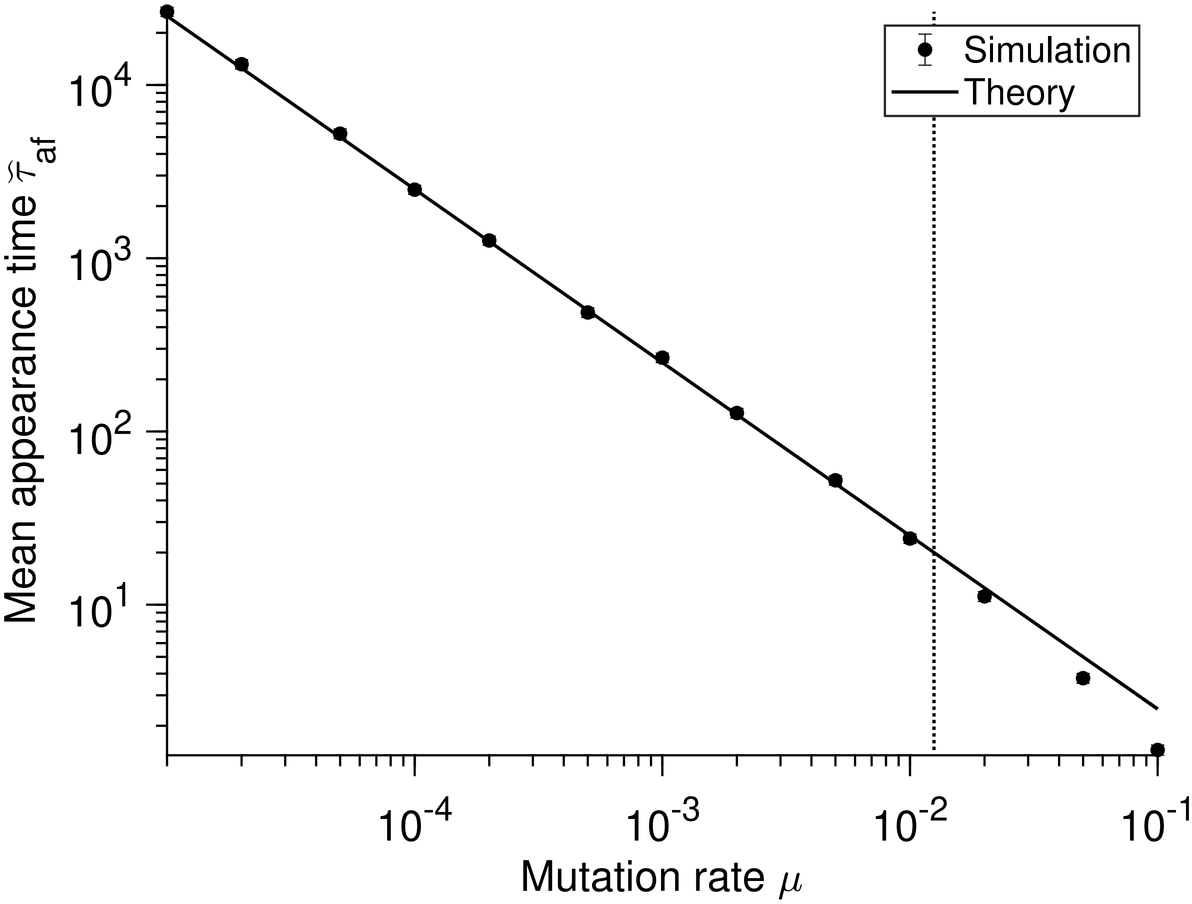


Figure 2: **The mean appearance time of a mutant is inversely proportional to the mutation rate in a constant environment.** Mean appearance time $\widetilde{\tau}_{\mathrm{af}}$ of mutation versus mutation rate $\mu$ in an effective environment resulting from fast environmental fluctuations. Each data point is averaged over $10^3$ stochastic realizations. The solid line corresponds to the analytical prediction $1/(\mu\, \widetilde{N}_W^*\, \widetilde{d}_W\, \widetilde{s})$. The Logistic model is used for this figure. Parameter values: $\widetilde{b}_W = 1$, $d_W = 0.1$, $b_M = 1$, $d_M$, $K = 100$, $\widetilde{N}_W^* = 80$ and $\widetilde{s} = 0.5$. For these parameters, $\widetilde{T}_0 \sim 10^{35}$.

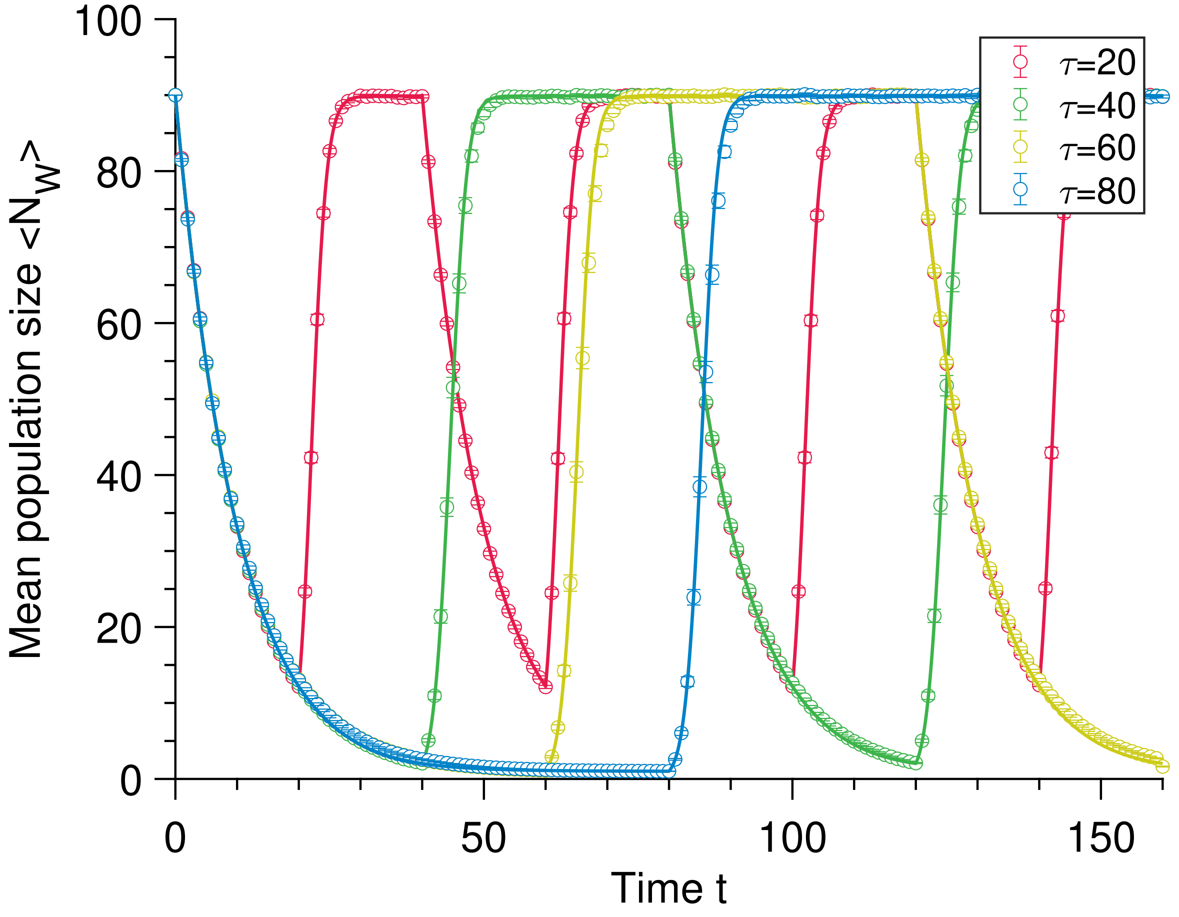


Figure 3: **The population size is periodic in a fluctuating environment.** Mean population size $\langle N_W \rangle$ versus time $t$ in a switching environment for different phase durations $\tau$. Each data point is averaged over $10^3$ stochastic realizations. The solid line corresponds to the analytical predictions (see main text). The Logistic model is used for this figure. Parameter values: $b_{W,F} = 1$, $b_{W,H} = 0$, $d_W = 0.1$, $K = 100$ and $N_W^* = 90$.

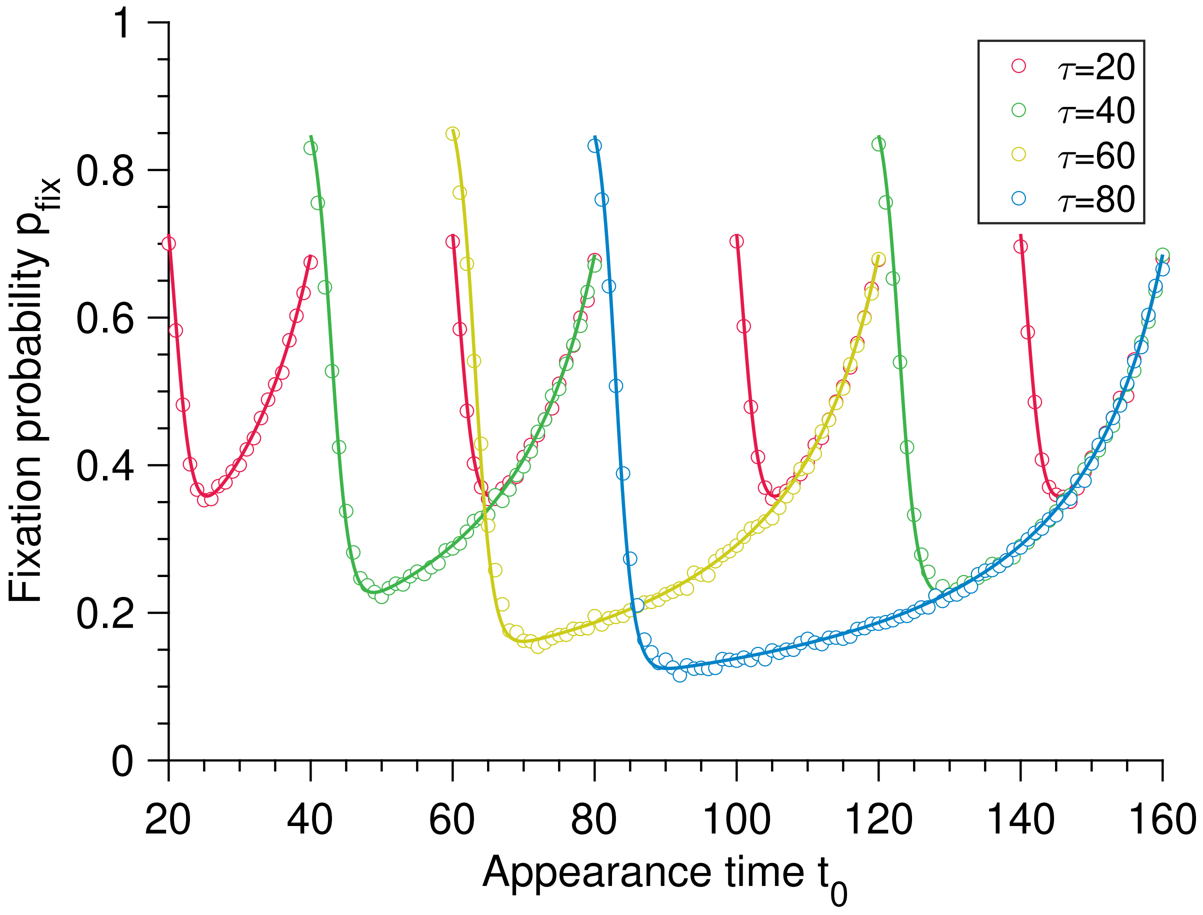


Figure 4: **The fixation probability is periodic in a fluctuating environment.** Fixation probability $p_{\text{fix}}$ of mutation versus appearance time $t_0$ in a fluctuating environment for different phase durations. Each data point is averaged over $10^3$ stochastic realizations. The solid line corresponds to the analytical predictions (see main text). The Logistic model is used for this figure. Parameter values: $b_{W,F} = 1$, $b_{W,H} = 0$, $d_W = 0.1$, $b_M = 1$, $d_M = 0.1$, $K = 100$ and $N_W^* = 90$. Note that no data is displayed in the phases where $b_{W,H} = 0$ since no division occurs.

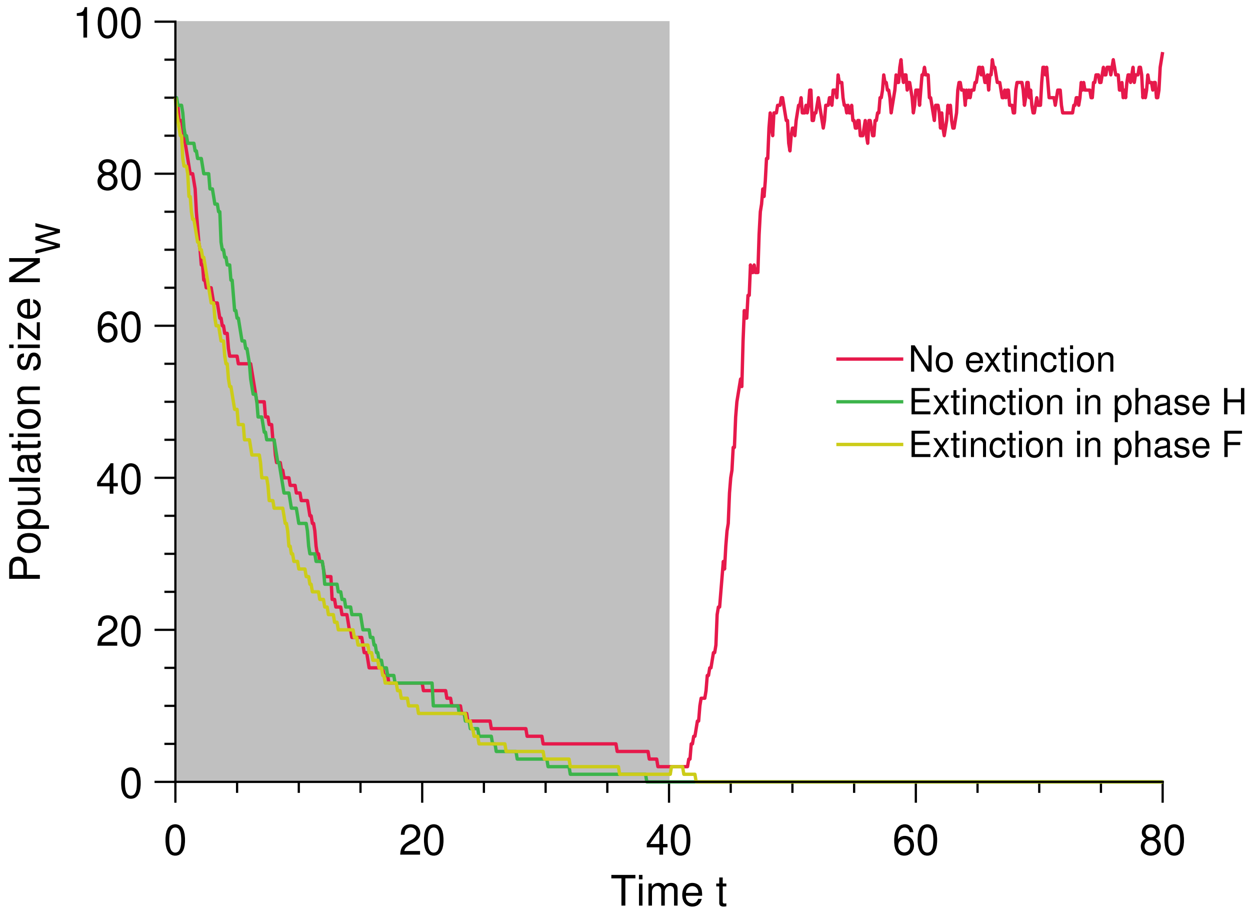


Figure 5: **Two different mechanisms can drive the population to extinction.** Population size $N_W$ versus time $t$ in a switching environment. Each line represents a stochastic realization with different outcomes. The gray and white phases correspond to harsh and favorable environments, respectively. The Logistic model is the model used for this figure. Parameter values: $b_{W,F} = 1$, $b_{W,H} = 0$, $d_W = 0.1$, $K = 100$ and $N_W^* = 90$.

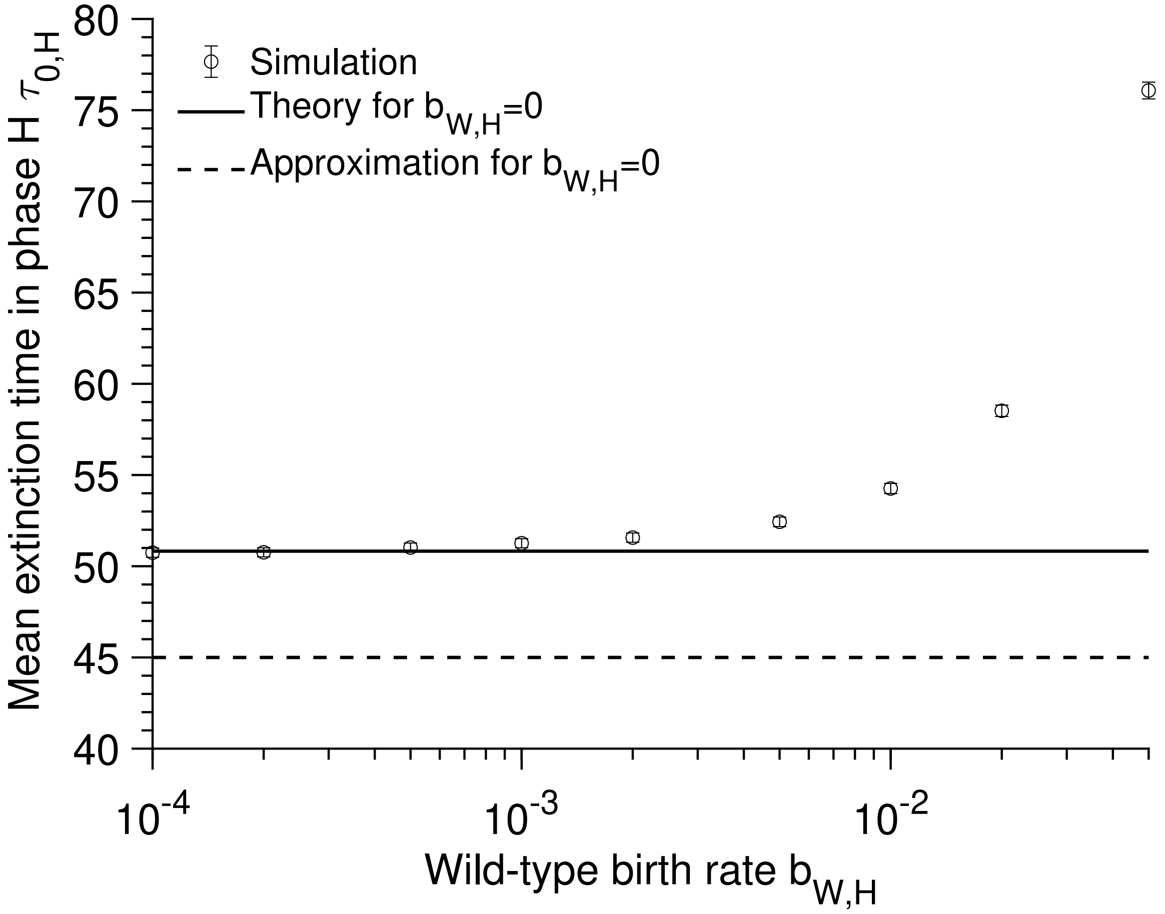


Figure 6: **Survival time in a deleterious environment.** Mean extinction time $\tau_{0,H}$ versus wild-type birth rate $b_{W,H}$ in the harsh environment. Each data point is averaged over $10^3$ stochastic realizations and displayed with its 95% confidence interval. The horizontal solid and dashed lines correspond to the equations $(1/d_W)\sum_{i=1}^{N_W^*} 1/i$ (exact for $b_{W,H} = 0$) and $\log(N_W^*)/d_W$ (approximation for $b_{W,H} = 0$), respectively. Parameter values: $d_W = 0.1$, $K = 100$ and $N_W^* = 90$.

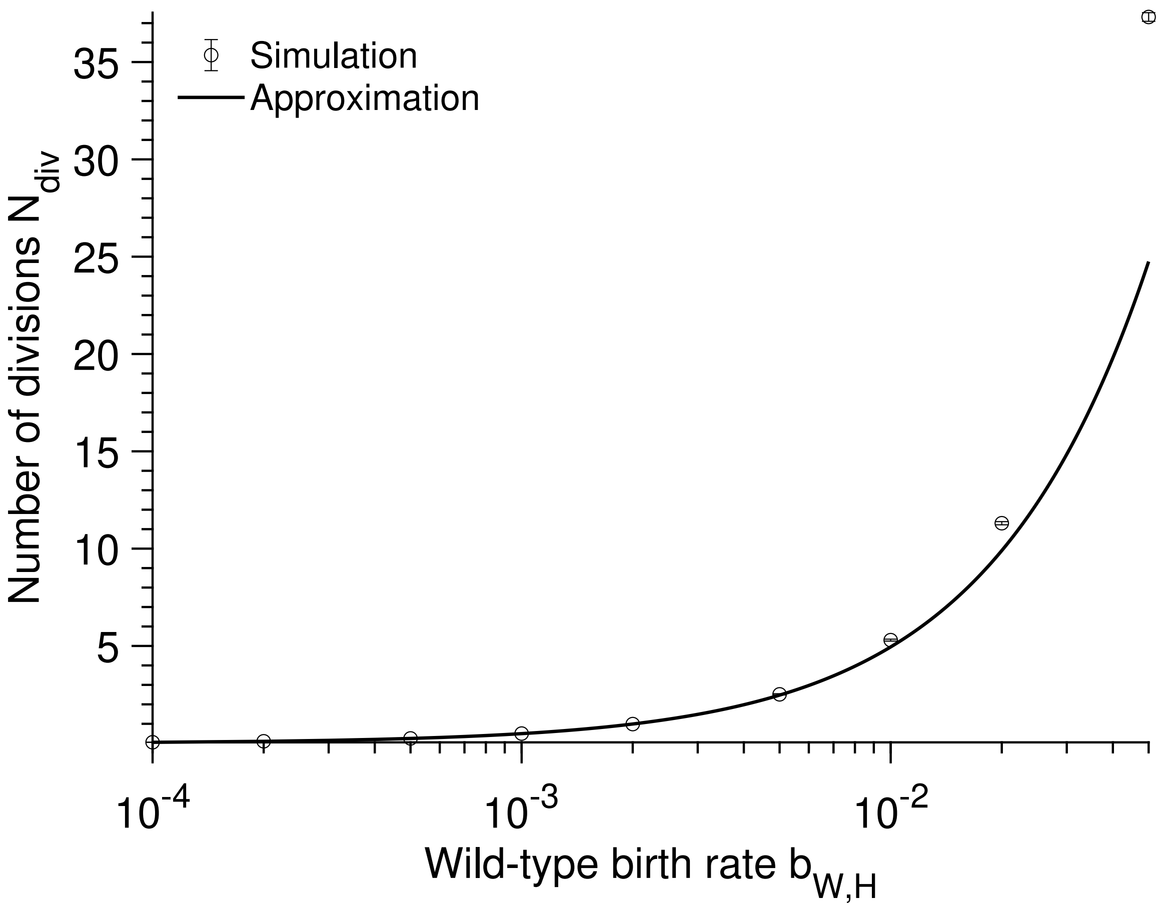


Figure 7: **There are few reproductions in the harsh environment when the birth rate is small.** Mean number of divisions $N_{\text{div}}$ versus wild-type birth rate $b_{W,H}$ in the harsh environment. Each data point is averaged over $10^3$ stochastic realizations and displayed with its 95% confidence interval. The solid line correspond to the equations $b_{W,H}\, N_W^*\, (2K - N_W^*)/(2 d_W\, K)$. Parameter values: $d_W = 0.1$, $K = 100$ and $N_W^* = 90$.

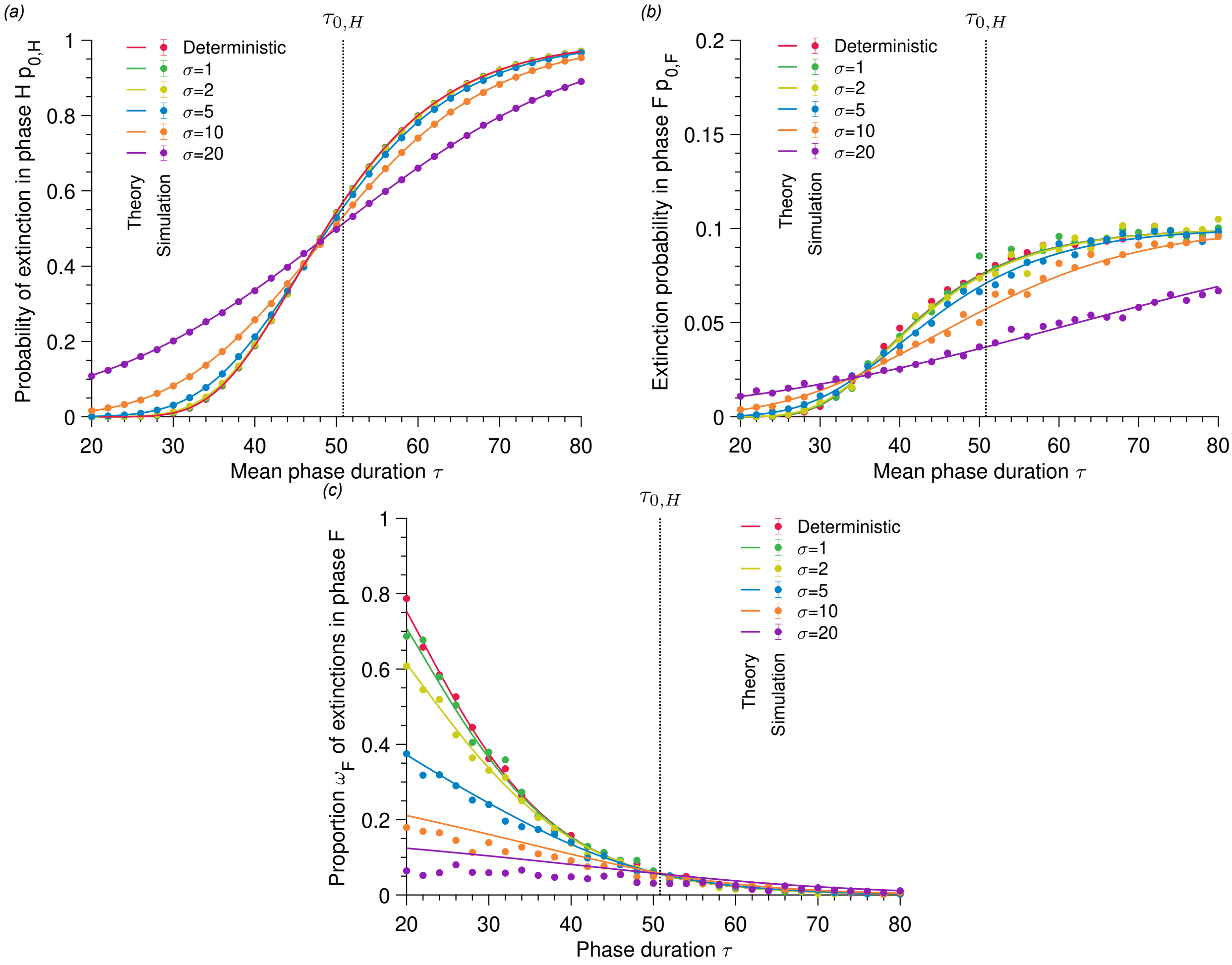


Figure 8: **The extinction probabilities increase with the phase duration.** *(a)* Probability of extinction in the harsh environment $p_{0,H}$ versus mean phase duration $\tau$ for deterministic and stochastic environmental fluctuations. *(b)* Probability of extinction in the favorable environment $p_{0,F}$ versus mean phase duration $\tau$ for deterministic and stochastic environmental fluctuations. *(c)* Proportion of extinction in the favorable environment $\omega_F$ versus mean phase duration $\tau$ for deterministic and stochastic environmental fluctuations. Each data point is averaged over $10^4$ stochastic realizations. The solid line corresponds to our analytical predictions (see main text). The dotted lines show the mean lifetime in the harsh environment. Parameter values: $b_{W,F} = 1$, $d_W = 0.1$, $b_{W,H} = 0$, $K = 100$ and $N_W^* = 90$.

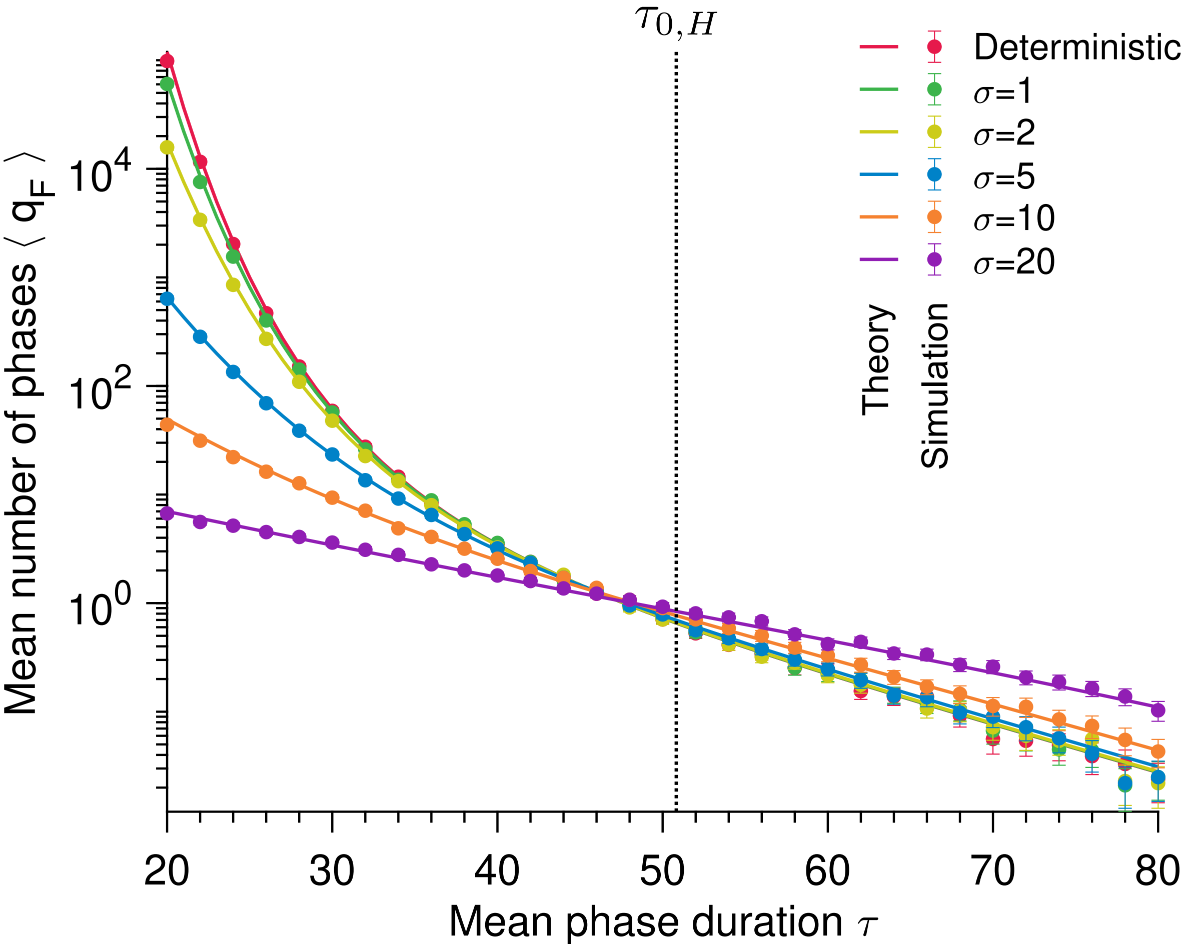


Figure 9: **The number of favorable phases decreases with the phase duration.** Mean number of favorable phases before extinction $\langle q_F \rangle$ versus mean phase duration $\tau$ for deterministic and stochastic switches. Each data point is averaged over $10^4$ stochastic realizations. The solid lines correspond to our analytical predictions (see main text). The dotted line shows the mean lifetime in the harsh environment. Parameter values: $b_{W,F} = 1$, $d_W = 0.1$, $b_{W,H} = 0$, $K = 100$ and $N_W^* = 90$.

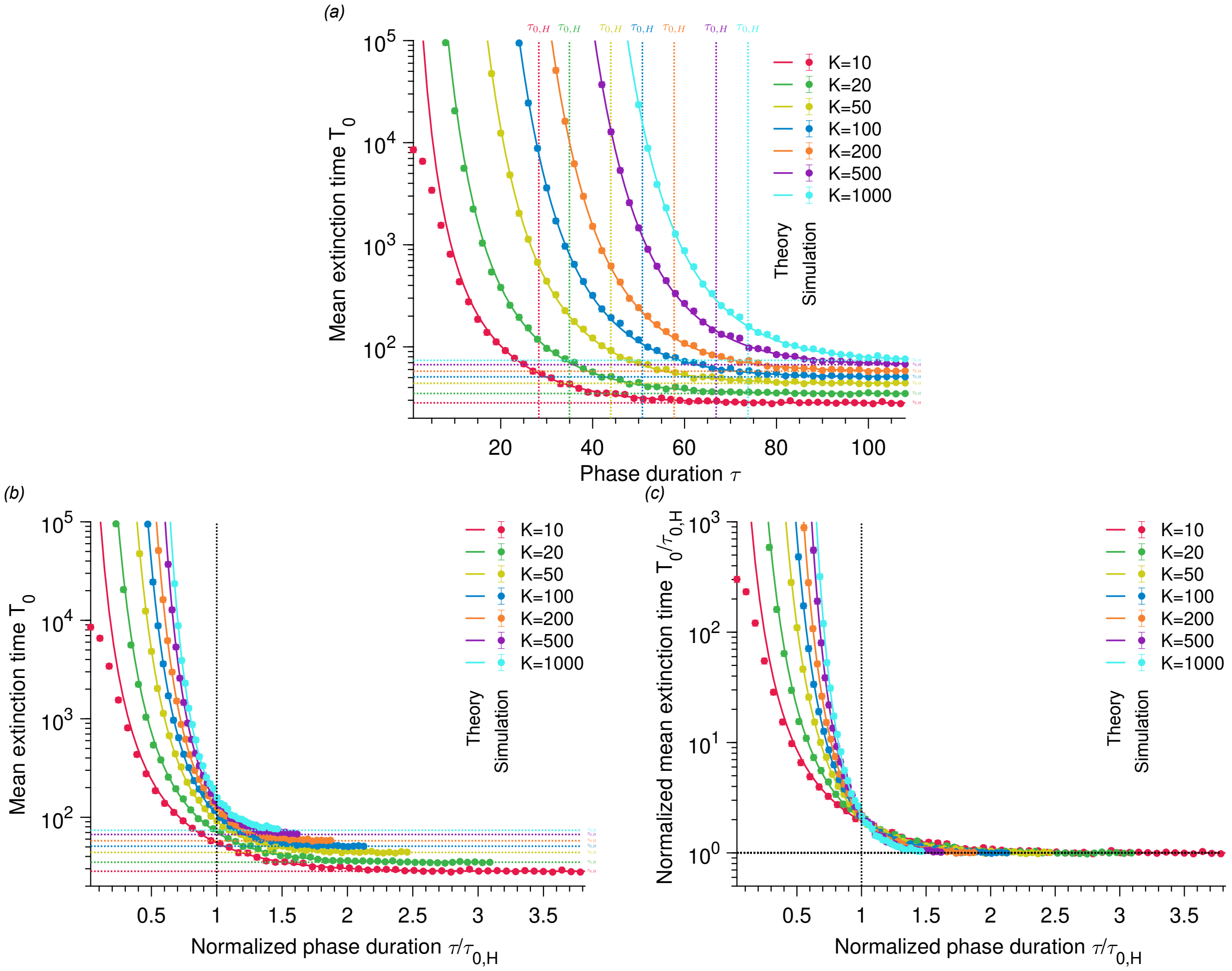


Figure 10: **The mean extinction time increases with the carrying capacity and decreases with the phase duration.** Mean extinction time $T_0$ versus phase duration $\tau$ for deterministic and different carrying capacities $K$. In *(b)*, the x-axis is normalized by the lifetime in the harsh environment. In *(c)*, both axes are normalized by the lifetime in the harsh environment. Each data point is averaged over $10^4$ stochastic realizations. The solid lines correspond to our analytical predictions (see main text). The dotted line shows the mean lifetime in the harsh environment. Parameter values: $b_{W,F} = 1$, $d_W = 0.1$, $b_{W,H} = 0$.

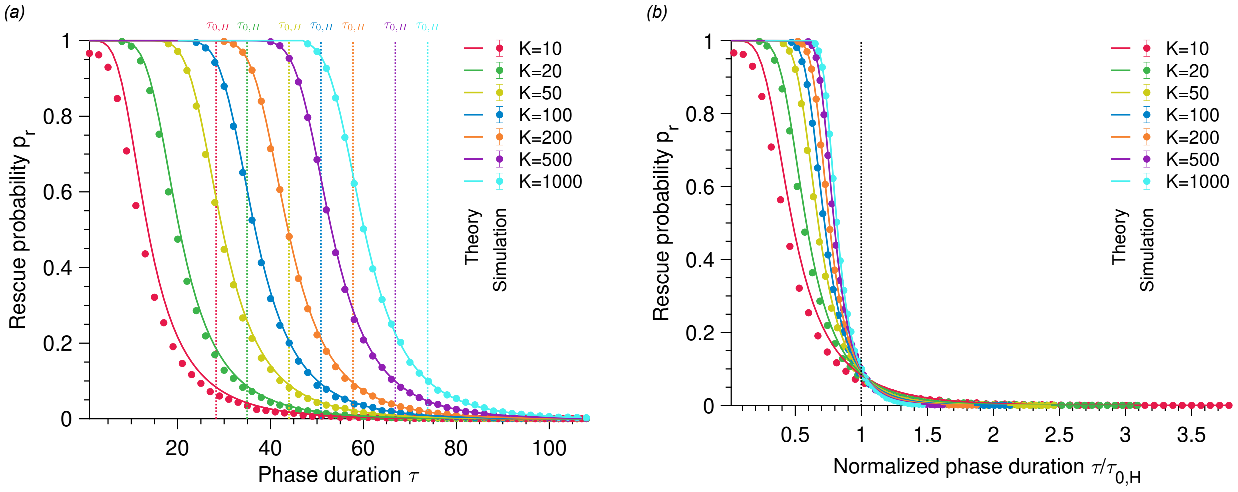


Figure 11: **The rescue probability is shifted towards increasing phase durations as the carrying capacity increases.** Rescue probability $p_r$ versus phase duration $\tau$ for deterministic and different carrying capacities $K$. In both panels, the product carrying capacity times mutation rate (i.e., $K\mu$) is the same for each data set. In *(b)*, the x-axis is normalized by the lifetime in the harsh environment. Each data point is averaged over $10^4$ stochastic realizations. The solid lines correspond to our analytical predictions (see main text). The dotted line shows the mean lifetime in the harsh environment. Parameter values: $b_{W,F} = 1$, $d_W = 0.1$, $b_{W,H} = 0$, $b_M = 1$, $d_M = 0.1$ and $\mu = 10^{-1}/K$.

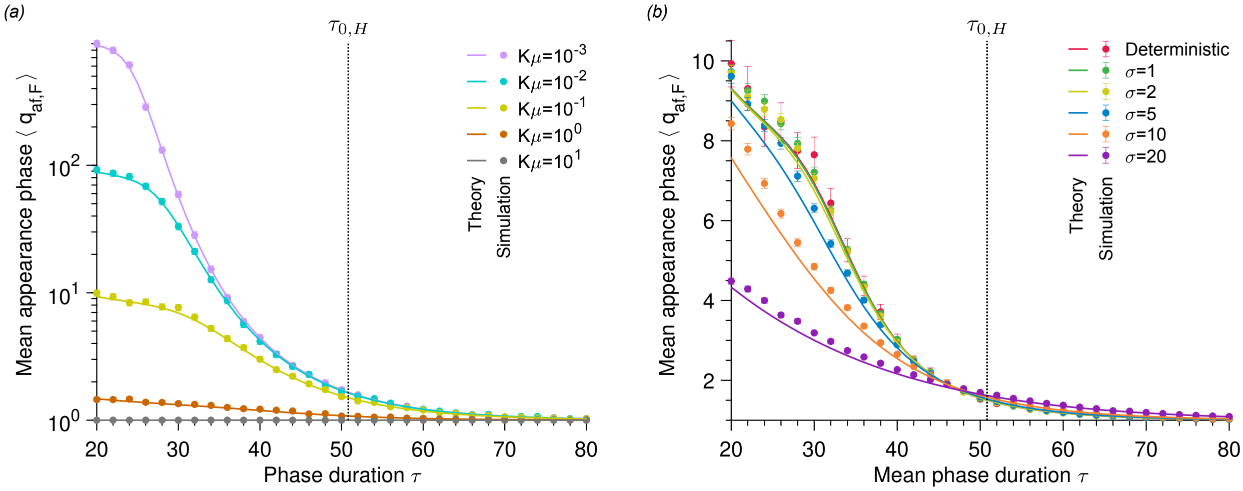


Figure 12: **The mean number of favorable phases before the appearance of a mutant that fixes decreases as the phase duration increases.** Mean number of phases F $q_{\mathrm{af},F}$ versus phase duration $\tau$ for different mutation rates $\mu$ *(a)* as well as deterministic and stochastic environmental fluctuations *(b)*. Each data point is averaged over $10^4$ stochastic realizations. The solid lines correspond to our analytical predictions (see main text). In *(a)*, the environmental fluctuations are deterministic. The dotted line shows the mean lifetime in phase H. Parameter values: $b_{W,F} = 1$, $d_W = 0.1$, $b_{W,H} = 0$, $K = 100$, $N_W^* = 90$ and $\mu = 10^{-3}$ (in $a$).
